# KLHDC7B as a Potential Therapeutic Target in Systemic Lupus Erythematosus: Insights from Bulk and Single-Cell RNA-Seq and Spatial Transcriptomics of Epidermal Keratinocytes

**DOI:** 10.1101/2024.07.24.604953

**Authors:** Sen Guo

## Abstract

Systemic lupus erythematosus (SLE) is an autoimmune disease in which the body’s immune system mistakenly attacks healthy tissue in many body parts, symptoms vary among people and may be mild to severe. Red rash on the face is the most common symptom, others include painful and swollen joints, fever, chest pain, and feeling tired [1]. Both genetics and environmental factors are identified to be involved in SLE [2]. Currently, there is no cure for SLE, and corticosteroids are usually used as one of the treatment methods, but long-term use leads to side effects [3]. Identifying new therapeutic targets is crucial for improving the treatment of Systemic Lupus Erythematosus (SLE). By integrating bulk and single-cell RNA-seq data with spatial transcriptomics of keratinocytes, we found KLHDC7B to be upregulated in UV-treated HaCaT cells, SLE skin keratinocytes, and spatially resolved epidermal samples. These findings suggest KLHDC7B as a potential therapeutic target for SLE.

## Introduction

Systemic lupus erythematosus (SLE) is the most prevalent form of lupus, affecting approximately 70% of individuals diagnosed with the disease. SLE can lead to acute or chronic inflammation of multiple organs or organ systems. In contrast, cutaneous lupus erythematosus (CLE) is confined to the skin, although it may progress to SLE in some patients. Drug-induced lupus erythematosus (DILE) results from certain prescription medications and shares many clinical manifestations with SLE but typically does not involve major organs and resolves within six months of discontinuing the causative medication. Neonatal lupus, which manifests exclusively in newborns, is not considered a true form of lupus. The symptoms of neonatal lupus generally resolve within six months.

Global prevalence rates of SLE range from approximately 20 to 70 per 100,000 individuals. Among females, the incidence peaks between the ages of 45 and 64 years. The lowest prevalence rates are observed in Iceland and Japan, while the highest rates are found in the United States and France. The reasons for these geographic variations are not well understood, and environmental factors may play a significant role. For instance, differential exposure to sunlight and ultraviolet (UV) radiation across countries could influence the dermatological manifestations of SLE [4].

Systemic lupus erythematosus (SLE) affects both males and females; however, it is significantly more prevalent in women [5]. The clinical manifestations of SLE exhibit sex-specific differences. In females, the disease is characterized by a higher frequency of relapses, leukopenia, arthritis, Raynaud’s phenomenon, and psychiatric symptoms. In contrast, males with SLE are more prone to experiencing seizures, renal involvement, serositis, dermatological manifestations, and peripheral neuropathy [6].

Systemic lupus erythematosus (SLE) demonstrates familial aggregation, although no single causative gene has been identified. Instead, multiple genes contribute to the susceptibility to SLE, interacting with environmental triggers. Genes within the human leukocyte antigen (HLA) class I, class II, and class III regions are associated with SLE, with classes I and II independently contributing to an increased risk of the disease [7]. Other genes implicated in SLE risk include IRF5, PTPN22, STAT4, CDKN1A, ITGAM, BLK, OX40L, and BANK1 [8].

Certain susceptibility genes may exhibit population-specific associations. Genetic studies support a heritable component to SLE, with a heritability estimate exceeding 66%. Monozygotic twins exhibit a concordance rate of over 35% for SLE, compared to 2–5% in dizygotic twins and other full siblings, underscoring the genetic basis of the disease [9].

Increased ultraviolet (UV) exposure has been correlated with a heightened risk of malar rash at presentation [10]. In this study, we first analyzed the public dataset GSE198792, which compares UVB-treated (50 mJ) HaCat cells to controls. We subsequently examined the significantly upregulated genes in the single-cell RNA sequencing (scRNA-seq) dataset GSE179633, and finally in the epidermis of SLE patients compared to controls in the spatial transcriptomics dataset GSE182825. Our findings suggest that KLHDC7B may serve as a potential therapeutic target for SLE.

## Methods

### Bulk RNA-seq

Raw data of samples for SLE and control from GSE198792 were downloaded from the GEO datasets using the fastq-dump command from the SRA toolkit (v3.0.1) [11]. fastp (v0.23.4) was used to trim the adaptors and filter out the low-quality reads [12]. Basic twopassMode of STAR (v2.7.10b) was used to do the mapping of the clean reads to the human reference genome (GRCh38) from the GENCODE website [13]. featureCounts from subread (v2.0.6) was used to count the gene counts, using the countReadPairs parameter [14]. DESeq2 (v1.42.1) was used to identify the differentially expressed gene using the likelihood-ratio test (LRT) [15].

### scRNA-seq

The output files from Cell Ranger were downloaded from dataset GSE179633. The Seurat package (v5.1.0) was employed to import the quantification files and perform downstream analysis on 5 SLE samples and 2 healthy control samples [16]. Quality control was conducted based on VlnPlot results for nFeature_RNA (200 to 7500) and mitochondrial gene percentage (below 20%). SingleR (v2.4.1) was used to predict cell types in each sample [17]. All 5 SLE samples were combined into a single SLE Seurat object, and the 2 healthy control samples were combined into a single Control Seurat object. These two Seurat objects were subsequently merged into a unified Seurat object.

Normalization was performed on the final combined Seurat object, followed by the identification of variable features using FindVariableFeatures, data scaling with ScaleData, and differential expression analysis with FindMarkers to identify differentially expressed genes between SLE and control samples. Additionally, dimensionality reduction techniques, including RunPCA, RunUMAP, and RunTSNE, were applied to the combined Seurat object to facilitate the visualization of specific genes of interest using FeaturePlot. Gene set enrichment analysis was applied to the DEGs by the GAGE R package using all the genes and the corresponding log2FC values. [18]

### Spatial transcriptome

Analyzed counts of the epidermis from the GSE182825 were downloaded and used DESeq2 (v1.42.1) to identify the differentially expressed genes.

## Results

Significant upregulated genes after adjusted pvalue < 0.05 and |log2FC| >1 of UVB treated vs. Control HaCaT samples were identified, including the inflammatory-related genes of IL6, ZMYND10-AS1, GAST, KLHDC7B, KIF17, ENHO, SPINK6, TCIM, STOM, INHBE, IL1R1, ADM2, FAHD2P1, ZBP1and MUC13 (Figure 1). GO term Regulation of interleukin-1-mediated signaling pathway by STRING database was identified (strength 2.75, FDR = 0.00058) [18]. The whole list of DEGs can be found in the Supplemental Table 1.

**Figure 1.**
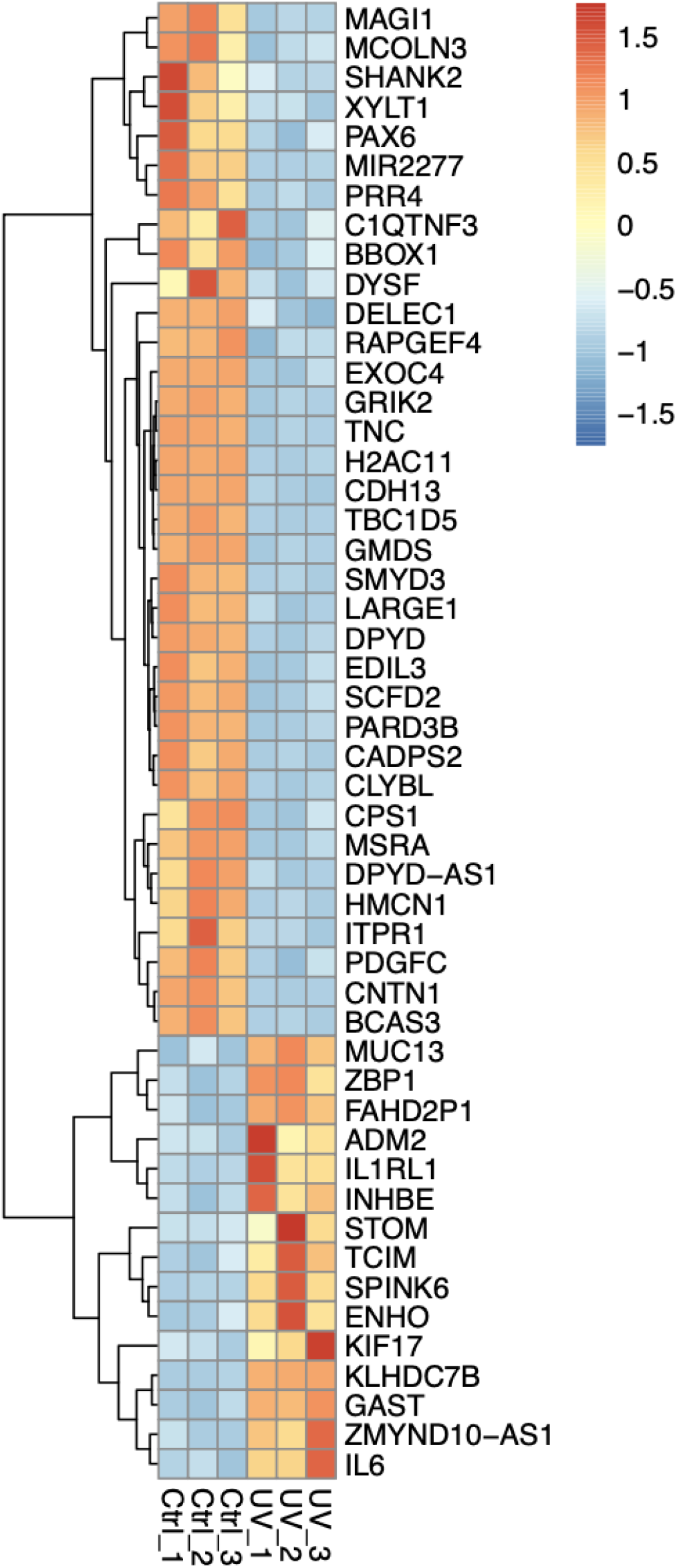
Significant differentially expressed genes of UVB treated vs. Control HaCaT samples

The singleR package predicts the presence of up to 20 distinct cell clusters within the dataset (Figure 2). However, the majority of the cells are classified as Keratinocytes. This observation aligns with the expected cellular composition of the epidermis, where Keratinocytes constitute the predominant cell type [19].

**Figure 2.**
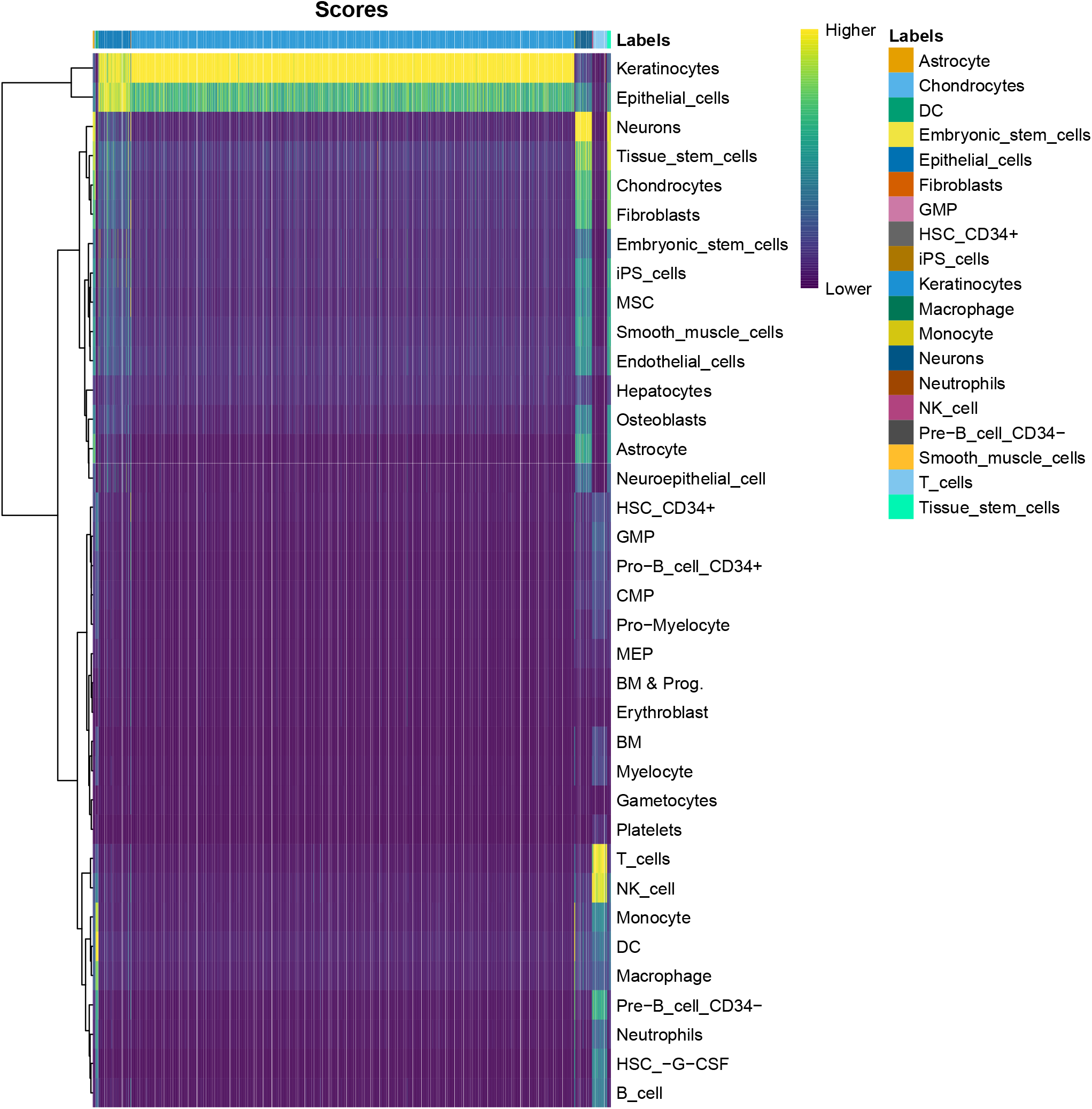
SingleR predictions for cell types in one healthy control sample (HC18) are shown, with yellow indicating the highest probability of cell type assignment.

The top differentially expressed genes (DEGs) under the threshold adjusted p value < 0.05 & |avg_log2FC| >1 between SLE and control samples are presented in the heatmap (Figure 3) and violin plot (Figure 4). The complete list of DEGs is available in Supplementary Table 2.

**Figure 3.**
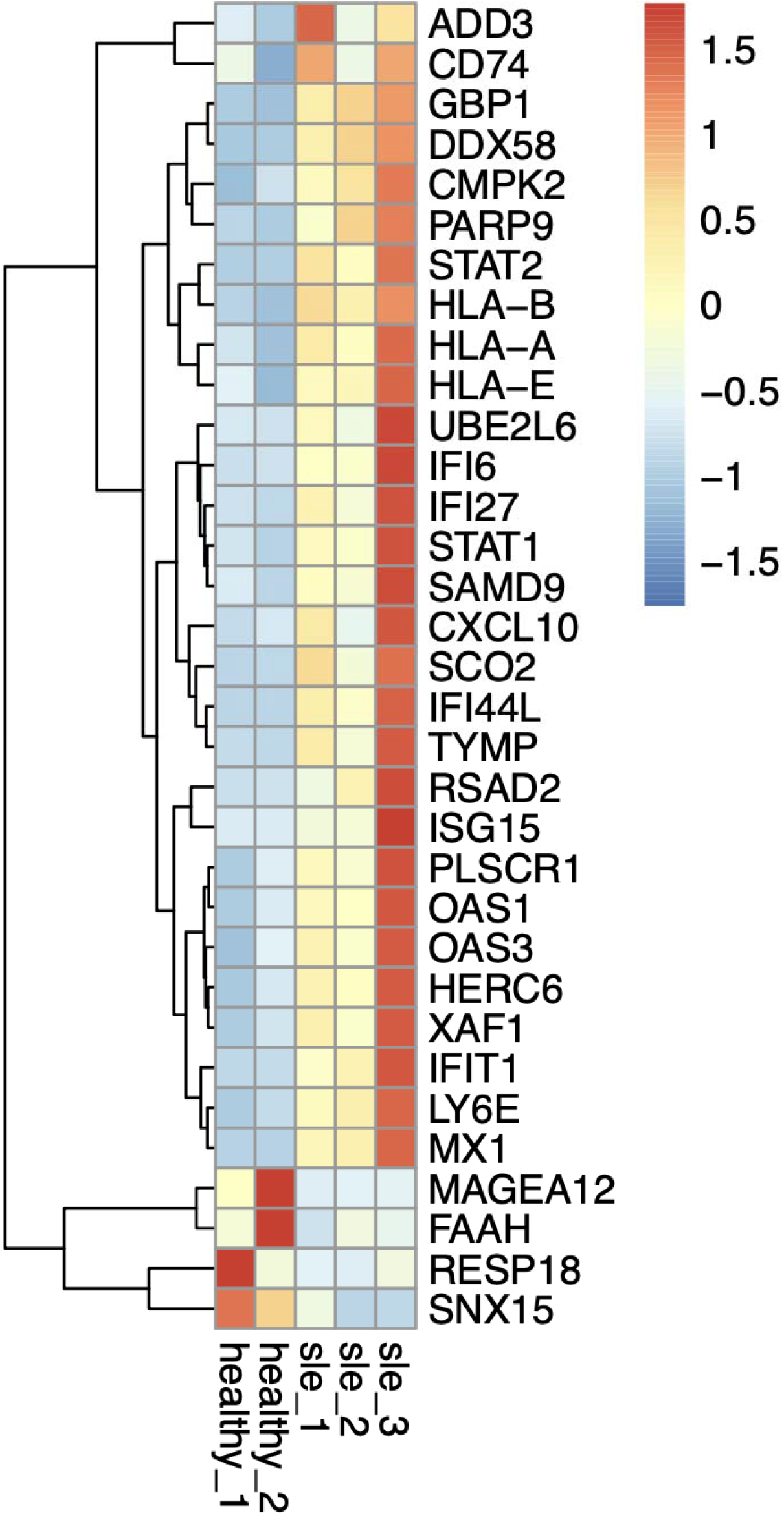
Top 33 significant differentially expressed genes of SLE vs. Control displayed.

**Figure 4.**
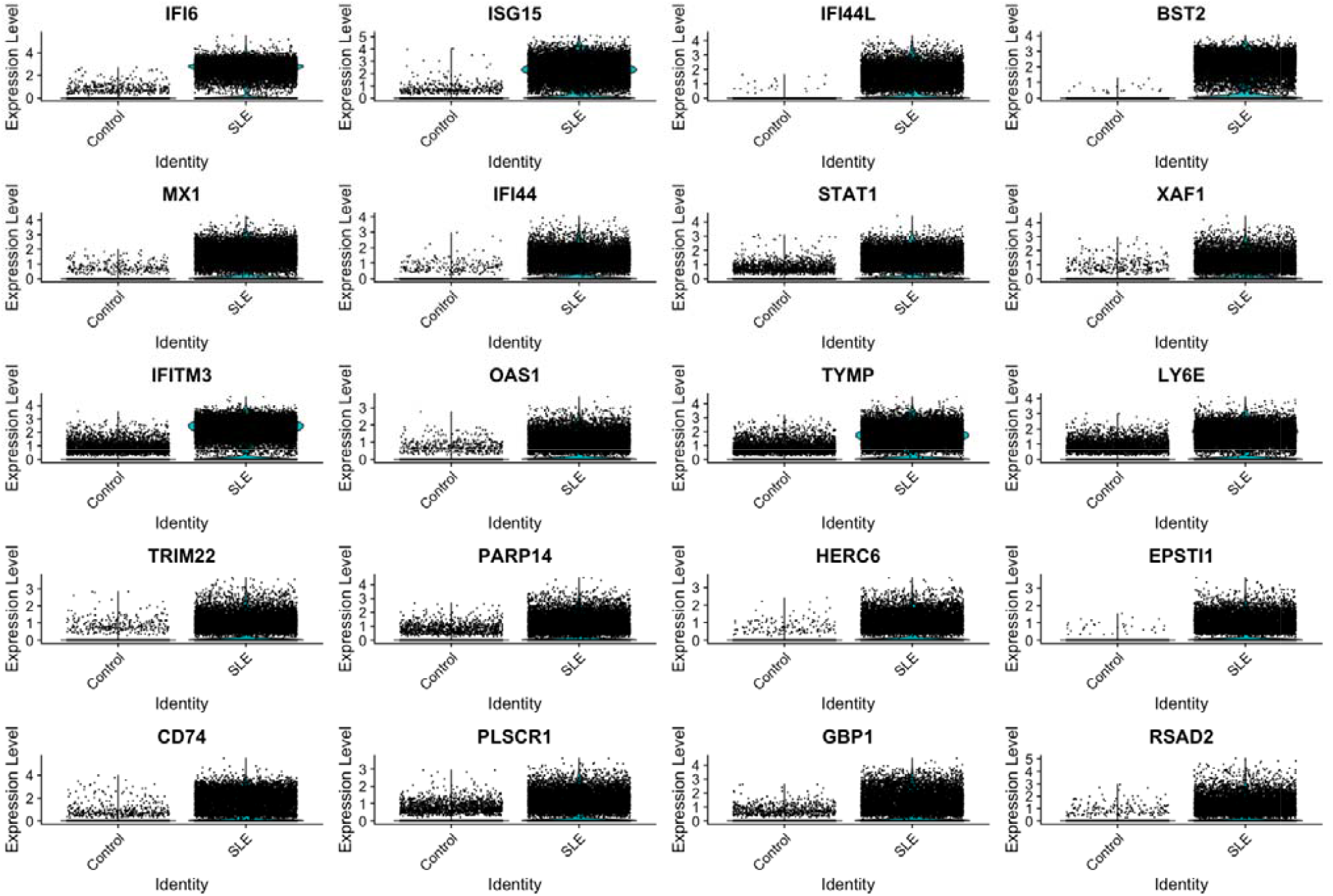
Violin plot of the top 20 differentially expressed genes in SLE and Control

Upon examining all significantly differentially expressed genes between UVB-treated and control HaCaT cells in the scRNA-seq data of SLE versus control samples, only KLHDC7B exhibited a distinct expression pattern. KLHDC7B was highly expressed in SLE cells but not control sample cells (Figure 5).

**Figure 5.**
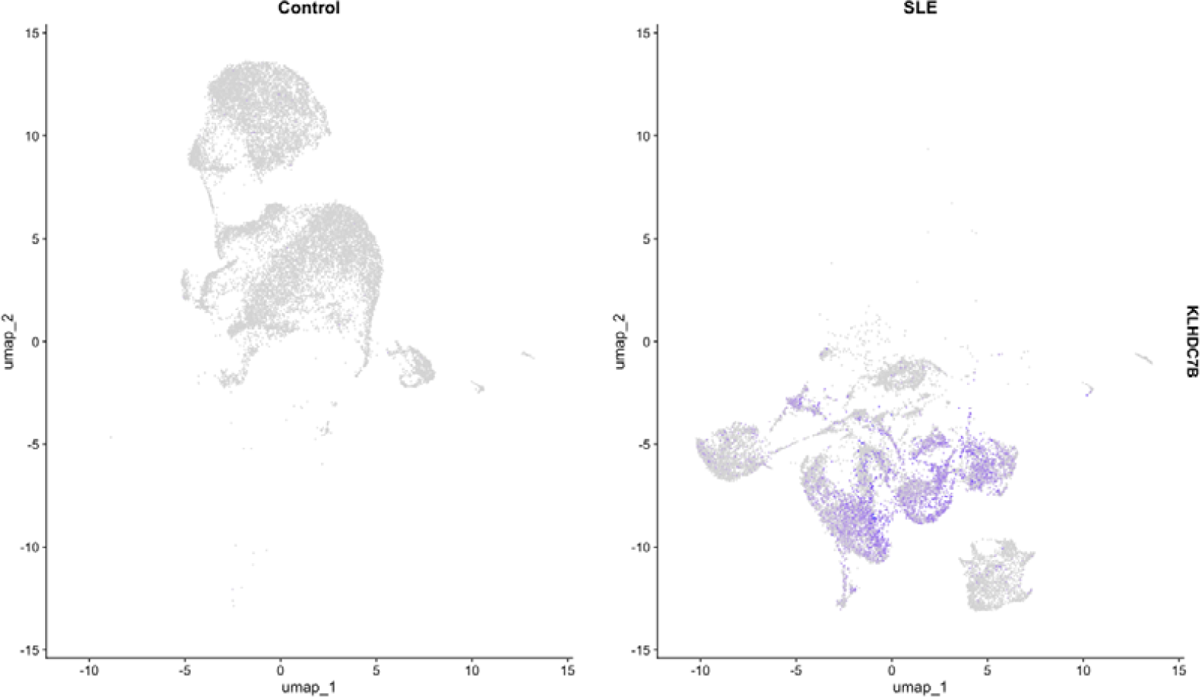
KLHDC7B Enrichment in SLE Sample Cells Compared to Control Sample Cells

Gene set enrichment analysis of differentially expressed genes in SLE versus control keratinocytes, derived from single-cell RNA sequencing data, revealed significant upregulation of interferon signaling pathways. Conversely, pathways associated with cell cycle regulation, translation, and other fundamental cellular processes were notably downregulated. These findings elucidate the molecular mechanisms potentially underlying the aberrant keratinocyte function in SLE pathogenesis.

**Figure 6.**
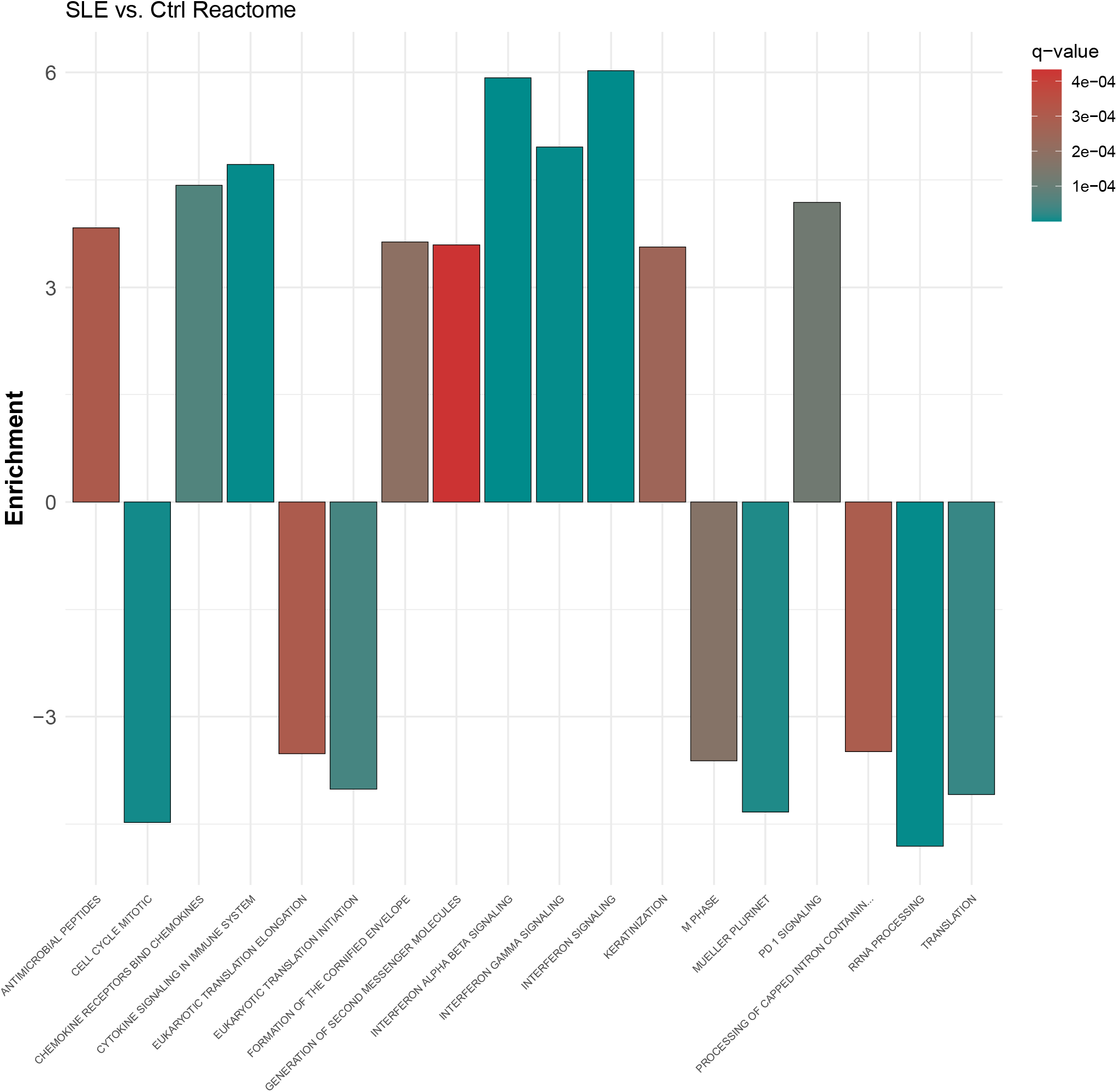
Gene set enrichment analysis of DEGs in SLE vs. Control of keratinocyte cells

The heatmap of top DEGs for the epidermis of SLE vs. Control samples is displayed in Figure 7 under the threshold p value < 0.01 & |log2FC| >1. Upregulated genes match the previous study, including the HLA-A/B/E, CXCL10, and ISG15 [20][21][22]. KLHDC7B shows the trend of upregulation with log2FC of 1.03. The complete list of differentially expressed genes (DEGs) is provided in Supplemental Table 3.

**Figure 7.**
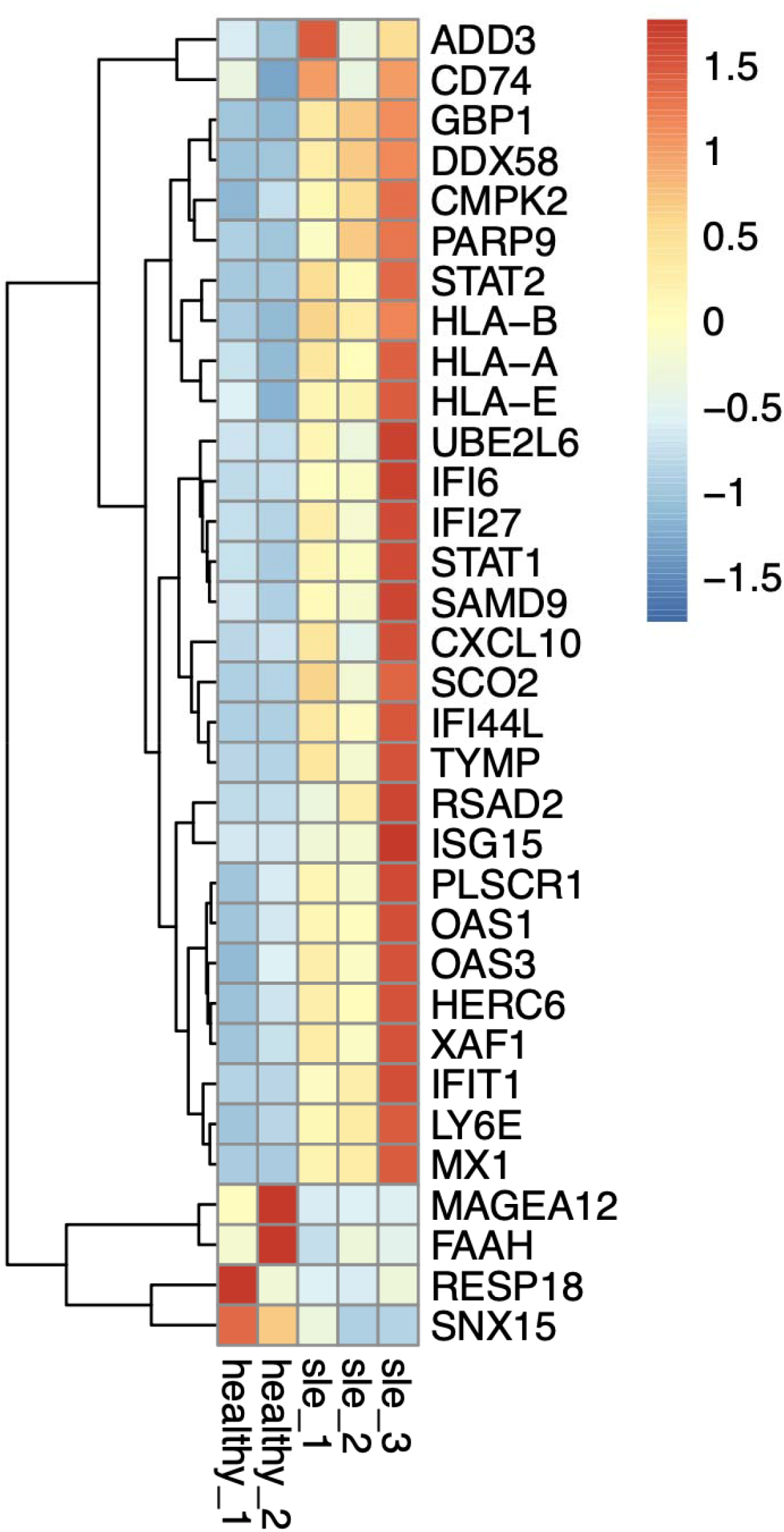
Top differentially expressed genes for the epidermis of SLE vs. Control samples

KLHDC7B contains the Kelch domain, there is study shows the drug Dimethyl fumarate targets Kelch-like proteins have pharmacological action [24] and applied to the SLE and can be a novel therapeutic agent to treat SLE [25].

## Conclusion

Systemic lupus erythematosus (SLE) is a complex and heterogeneous autoimmune disease that can affect multiple organ systems, including the skin, joints, kidneys, and nervous system. Despite significant advances in understanding the disease, the precise mechanisms underlying SLE pathogenesis remain incompletely understood. Recent integrative analyses of diverse data sets have identified KLHDC7B as a potential therapeutic target. This discovery opens new avenues for research into the molecular pathways involved in SLE and highlights the potential for novel therapeutic strategies aimed at modulating KLHDC7B activity.

KLHDC7B (Kelch Domain Containing 7B) has emerged as a candidate gene of interest. This gene is implicated in various cellular processes, including ubiquitination and protein degradation, which are critical in maintaining cellular homeostasis. Dysregulation of these processes is known to contribute to autoimmune disorders. Studies have shown that KLHDC7B expression levels are altered in SLE patients, suggesting its involvement in disease pathogenesis.

Further research is needed to elucidate the exact role of KLHDC7B in SLE. Investigations into how this gene interacts with other cellular pathways and contributes to immune system dysfunction will be essential. Additionally, exploring the therapeutic potential of targeting KLHDC7B could lead to the development of new treatments that better manage or even prevent disease progression.

In summary, while SLE remains a challenging disease with many unresolved questions, the identification of KLHDC7B as a potential therapeutic marker represents a promising step forward. Continued research efforts are crucial to translating these findings into clinical practice, ultimately improving outcomes for patients with SLE.

